# p53 knockout drives stage-specific long non-coding RNA reprogramming during human neural differentiation and glioma organoid tumorigenesis

**DOI:** 10.64898/2026.09.17.752309

**Authors:** Fabio Stasi, Alireza Rajabzadeh Manzari, Velia Siciliano, Elena Binda, Valerio Pazienza, Virgilio Brunetti, Massimo De Vittorio

**Author notes:** **Corresponding Authors:** Fabio Stasi, Center for Biomolecular Nanotechnologies, Istituto Italiano di Tecnologia, Arnesano (LE), Italy, Alireza Rajabzadeh Manzari, Center for Biomolecular Nanotechnologies, Istituto Italiano di Tecnologia, Arnesano (LE), Italy. Fabio Stasi and Alireza Rajabzadeh Manzari contributed equally to this work and share co-first authorship.

## Abstract

**Background:** Long non-coding RNAs (lncRNAs) are emerging regulators of tumor initiation and progression through their effects on stemness, lineage commitment, and transcriptional plasticity. The contribution of lncRNA reprogramming during early p53-deficient gliomagenesis remains poorly understood.

**Methods:** Human induced pluripotent stem cells (iPSCs) carrying a complete TP53 knockout were differentiated into neural progenitor cells (NPCs) and cerebral organoids. Bulk RNA sequencing was performed at multiple differentiation stages and compared with established glioblastoma cell lines (U87 and U118) to identify lncRNA expression changes associated with p53 loss and tumor-like transformation.

**Results:** p53 deficiency induced a progressive and stage-specific remodeling of the lncRNA landscape, with the most pronounced transcriptional shift observed in 40-day NPCs. This late-stage profile partially overlapped with glioblastoma-like cell lines and was characterized by the upregulation of five lncRNAs enriched in both p53-deficient neural models and glioma cells. Principal component analysis showed that these lncRNAs clustered with tumor-associated samples and were strongly co-expressed with mitotic cell-cycle genes. In particular, LINC00973 and LINC01583 were highly expressed in primary glioblastoma datasets, while RP5-875H18.9 and RP11-1094H24.4 emerged as previously unrecognized glioma-associated transcripts.

**Conclusion:** Loss of p53 drives stage-dependent lncRNA reprogramming during neural differentiation and promotes the emergence of glioma-associated transcriptional features. These findings identify candidate lncRNAs that may contribute to early gliomagenesis and represent potential biomarkers or therapeutic targets in glioblastoma.

**Importance of the Study:** TP53 alterations are among the most common events in glioma, yet the transcriptional changes that accompany the earliest stages of tumor development remain incompletely understood. Using an isogenic human model based on TP53-knockout iPSCs, neural progenitor cells, and cerebral organoids, we identified a stage-specific transcriptional shift that emerges during late neural differentiation and resembles key features of glioblastoma. This transition is accompanied by the activation of a distinct lncRNA program, including previously unreported glioma-associated transcripts. The relevance of these findings is supported by independent validation in TCGA patient cohorts. Our study provides a human developmental framework to investigate the molecular events linking p53 loss to gliomagenesis and highlights novel non-coding RNA candidates for future mechanistic and translational studies.

**Key Points:** *Question:* How does p53 deficiency affect lncRNA expression during human neural differentiation?

*Findings:* We identified a late transcriptional program that emerges in p53-deficient neural progenitors and organoids and partially overlaps with glioblastoma cell lines. This program includes five lncRNAs co-expressed with mitotic genes, two of which have not previously been linked to glioma.

*Relevance:* These findings provide insight into the molecular changes that accompany p53 loss during early gliomagenesis and highlight novel non-coding RNA candidates for future functional studies.

**Highlights:**

- p53 deficiency triggers a late-onset lncRNA reprogramming specifically at the 40-day NPC stage.
- Five lncRNAs (LINC00973, LINC01583, RP11-1094H24.4, RP11-462L8.1, RP5-875H18.9) are co-induced with mitotic genes in p53-KO and glioma cells.
- RP5-875H18.9 and RP11-1094H24.4 are novel candidates not previously reported in glioma.
- LINC01583 high expression significantly correlates with poorer overall survival in TCGA-GBM patients.
- The iPSC–NPC–organoid model faithfully recapitulates the transcriptional landscape of primary GBM.

**Graphical Abstract:** 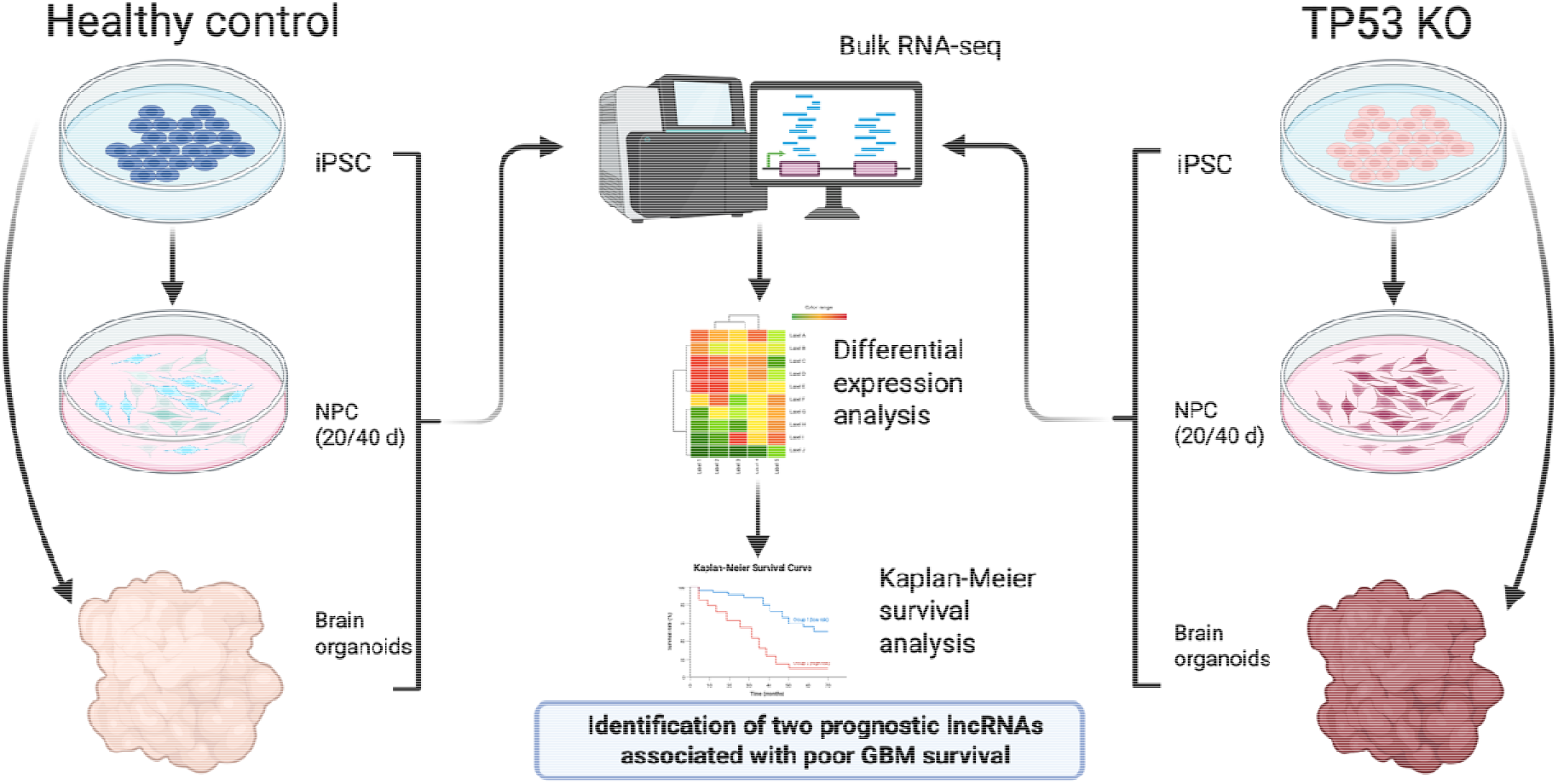

Comparative bulk RNA-seq analysis of human control and TP53 knockout iPSC- derived neural progenitor cells and brain organoids identified stage-specific lncRNA reprogramming during neural differentiation. Integration with glioblastoma patient survival data revealed two prognostic lncRNAs associated with poor clinical outcome.

## 1. Introduction

Glioblastoma (GBM) is an aggressive tumor of the central nervous system, characterized by high heterogeneity, significant intrinsic plasticity, and resistance to standard treatments [1]. Glioblastoma signifies a major failure in the cellular developmental hierarchy, with disruption of the p53 signaling pathway serving as a key regulator of neural stem cell (NSC) homeostasis. Genomic studies have shown alterations in the p53 pathway, with direct TP53 mutations present in 65% of secondary glioblastomas but in fewer than 30% of primary glioblastomas [2]. p53 is known as a barrier to dedifferentiation, and its impairment promotes conditions that keep cells in a plastic, undifferentiated state by diverting neural progenitor developmental pathways [3].

Long noncoding RNAs (lncRNAs) are transcripts longer than 200 nucleotides that have shown an emerging role in regulating tumor plasticity. These lncRNAs influence phenotypic transitions by acting as epigenetic modulators, transcriptional scaffolds, and competitive endogenous RNA (ceRNA) sponges. Several lncRNAs indirectly impact GBM progression, primarily through modulating miRNAs [4]. For instance, LINC00998, which interacts with CBX3, inhibits GBM cell proliferation via the c-Met/AKT/mTOR pathway and enhances patient survival [5]. Similarly, MDHDH serves as a scaffold for MDH2 and PSMA1, regulating NAD+ metabolism and autophagy to suppress tumor growth [6]. Additionally, lncRNAs play a key role in controlling stem-like transition and dedifferentiation. Recently, Pandey et al. discovered a notable increase in LINC00062 in lung adenocarcinoma cells, which is necessary for glucose-starvation-induced dedifferentiation [7].

In the context of GBM, despite clear evidence of interactions between lncRNA transcripts and epigenetically mediated miRNA modulations along with oncogenic signaling pathways (e.g., PI3K/AKT, Wnt/beta-catenin axes) [4], the mechanisms of co-evolution of coding and noncoding transcripts during the early stages of neural differentiation after p53 loss in controlled human models remain poorly understood. The temporal dynamics of this molecular reprogramming can be accurately mapped using human induced pluripotent stem cells (iPSCs) and 3D brain organoids, revealing strategic deficiencies in the tumor regulatory architecture [8]. Targeting the p53-lncRNA axis is a promising approach to disrupt the regulatory circuits that support glioblastoma survival and invasion.

An important question that remains underexplored is whether p53 deficiency triggers a distinct lncRNA reprogramming during GBM progression and the development of a malignant phenotype. So far, this process has not been thoroughly investigated or genomically mapped in detail. To address this research gap, a parallel isogenic human model system was designed using iPSCs carrying a complete p53 gene deletion (p53−/−). These cells differentiated into neural progenitor cells (NPCs) at 20 and 40 days and into brain tumor organoids in a separate study arm. Using an integrated approach based on total RNA sequencing (Total RNA-seq), RT-qPCR analysis, principal component analysis (PCA), and gene ontology enrichment (GO Enrichment), the question was raised whether the lack of p53 drives transcriptional changes in late phases of development and convergence towards a glioblastoma-like state.

## 2. Materials and Methods

### 2.1. Cell lines and culture conditions

Human glioblastoma cell lines U-87 MG (HTB-14) and U-118 MG (HTB-15) were obtained from the American Type Culture Collection (ATCC). Cells were maintained in DMEM high glucose supplemented with 10% fetal bovine serum (FBS), 1× non-essential amino acids, and 1× glutamine–alanine supplement, under standard culture conditions at 37°C in a humidified atmosphere with 5% CO□.

Human induced pluripotent stem cells (iPSCs) derived from a healthy donor (STEMCELL Technologies, SCTi005-A Healthy Control) were cultured in mTeSR Plus medium with supplement on plates coated with Matrigel matrix, hESC-qualified, according to the manufacturer’s recommendations.

### 2.2. Neural progenitor differentiation and organoid generation

Twenty-four hours after plating, iPSCs were induced toward the neural lineage using STEMdiff Neural Progenitor Basal Medium according to the manufacturer’s instructions. Medium was changed every 3 days, and cells were collected at 20 days and 40 days of differentiation for downstream analyses.

For organoid generation, iPSCs were differentiated using the STEMdiff Cerebral Organoid Kit according to the manufacturer’s protocol. Organoids were cultured for more than 40 days prior to analysis.

### 2.3. CRISPR/Cas9-mediated TP53 knockout

A TP53 knockout was generated using a CRISPR/Cas9 vector from VectorBuilder targeting exon 4 of the human TP53 locus. The guide-related oligonucleotide sequences used in the construct were CACCGCCATTGTTCAATATCGTCCG and AAACCGGACGATATTGAACAATGGC. Edited cells were expanded and used for downstream differentiation and molecular analyses.

### 2.4. RNA extraction, cDNA synthesis, and RT-qPCR

Total RNA was extracted from cultured cells and organoids using standard procedures. cDNA was synthesized from total RNA using SuperScript IV VILO Master Mix (Thermo Fisher Scientific) according to the manufacturer’s instructions. Quantitative real-time PCR was performed using TaqMan Gene Expression Assays on a QuantStudio 1 Real-Time PCR System (Thermo Fisher Scientific). For each reaction, 1 ng of cDNA was used as input template. Relative expression levels were calculated using the ΔΔCt method, with 18S rRNA used as the endogenous control for normalization.

The following TaqMan assays were used: SOX1 (Hs01057642_s1), PAX6 (Hs01088114_m1), TUBB3 (Hs00801390_s1), EGFR (Hs01076090_m1), Nestin (Hs04187831_g1), SOX2 (Hs04234836_s1), MAP2 (Hs00258900_m1), and 18S rRNA (Hs03003631_g1). Statistical significance across groups was assessed using ordinary one-way ANOVA followed by Tukey’s multiple-comparisons test in GraphPad Prism. P values are indicated in the corresponding figure panels.

### 2.5. RNA-seq analysis

RNA-seq data generated from healthy and TP53-knockout iPSCs, NPCs, and organoids were analyzed to identify coding and non-coding transcriptional changes across neural differentiation stages. Gene expression matrices were annotated using gene symbols and gene biotypes, and both protein-coding genes and non-coding RNAs were evaluated. Exploratory analyses included principal component analysis (PCA), fold-change-based prioritization across biologically relevant contrasts, identification of late KO / tumor-like transcriptional programs, and functional enrichment analysis using Gene Ontology Biological Process (GO- BP) and Reactome annotations.

### 2.6. lncRNA prioritization and public dataset validation

lncRNA candidates were prioritized based on enrichment in late TP53-deficient stages and tumor-like samples. Public glioma RNA-seq datasets from TCGA-GBM and TCGA-LGG were interrogated using TCGAbiolinks to assess external support for prioritized candidates. Expression differences between groups were evaluated using the Wilcoxon rank-sum test.

### 2.7. Bioinformatic analysis and visualization

Bioinformatic analyses and data visualization were performed in R, using packages including ggplot2, pheatmap, clusterProfiler, ReactomePA, and TCGAbiolinks.

## 3. Results

### 3.1. Differentiation of iPSCs into late neural progenitor cells and tumor organoids

To confirm neural differentiation and assess the effects of p53 deficiency on lineage development, the expression profile of marker genes in RNA-seq data was first analyzed. Pluripotency markers, including POU5F1/OCT4 and NANOG, were highly expressed in iPSC cells and rapidly downregulated upon differentiation. In contrast, early neural induction markers (PAX6, SOX1, and NES) peaked at day 20, while late neural maturation markers (TUBB3/TUJ1, MAP2, and DCX) reached maximum transcriptional levels at day 40 and were maintained in organoid models (Fig. S1). The validity of these transcriptional profiles was independently confirmed by RT-qPCR analysis on the same samples. The results showed that progenitor-associated markers (PAX6, SOX1, SOX2, and Nestin) were stably maintained or enriched in late p53-KO NPC. Conversely, the expression of neural differentiation-associated markers (TUBB3 and MAP2) was preserved or increased, particularly in p53-KO organoids. The expression of tumor-associated marker EGFR was significantly higher in p53-KO cells at late p53-KO NPC. Overall, the data confirm a successful shift to the late neural differentiation stage and reveal the context-dependent lineage remodeling induced by p53 deficiency: late NPC cultures preserve progenitor- and tumor-related features, while KO organoids retain features associated with mature neurons (Fig. 1).

**Fig. 1.**
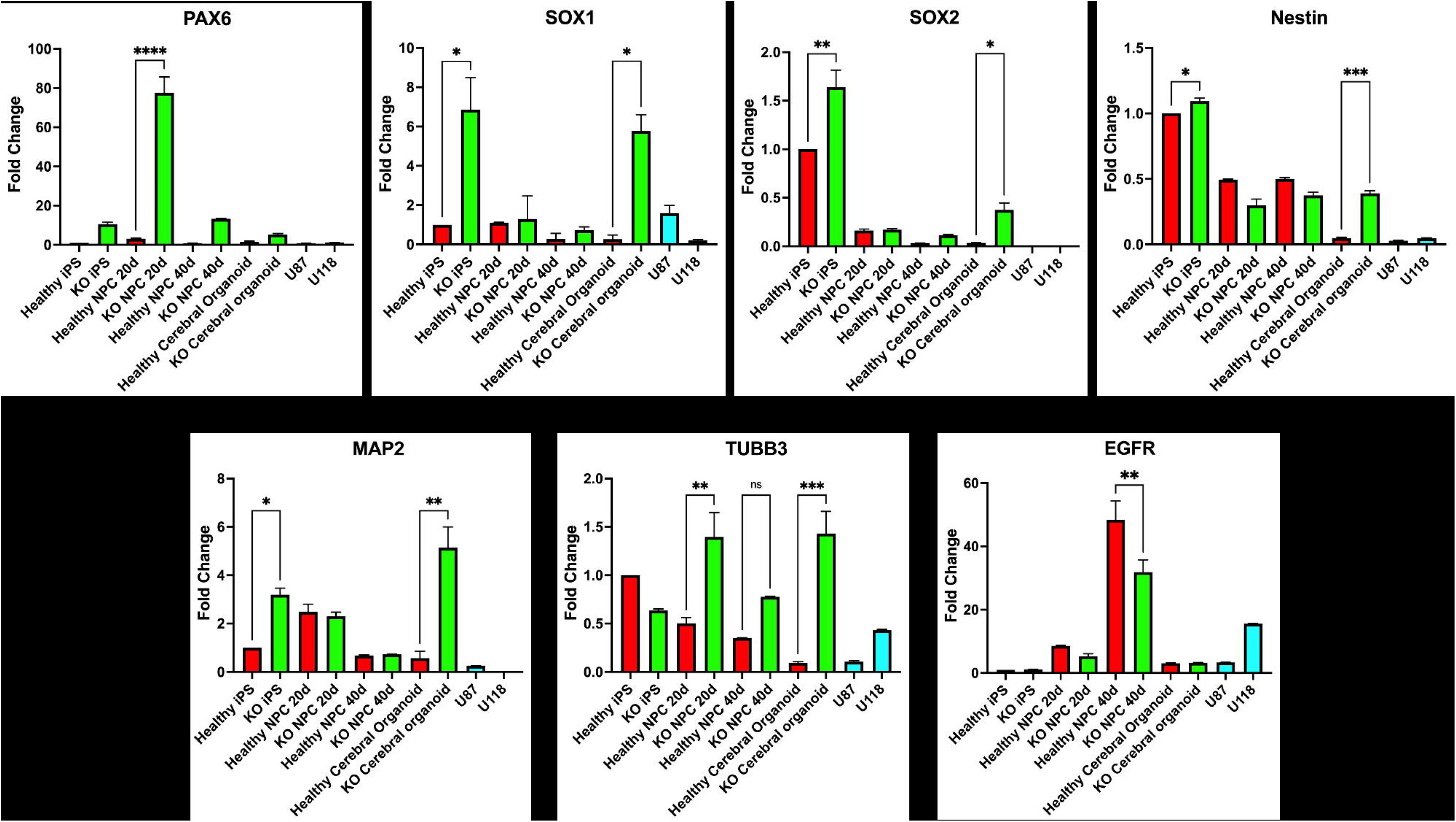
Relative fold-change expression (mean ± SEM) of progenitor-associated markers (PAX6, SOX1, SOX2, Nestin), neuronal differentiation markers (MAP2, TUBB3), and the tumor-associated marker EGFR, measured by RT-qPCR across healthy and p53-KO iPSCs, NPCs (20 d and 40 d), cerebral organoids, and glioma lines U87/U118. Statistical significance is indicated (*P < 0.05, **P < 0.01, ***P < 0.001, ****P < 0.0001; ns = not significant; one-way ANOVA with Tukey’s post-hoc test).

### 3.2. Prioritization of tumor-specific lncRNA candidates

To identify novel p53-loss-induced lncRNAs in the staged human iPSC–NPC–organoid model, we examined the late-stage tumor-like molecular signature. Five lncRNAs were prioritized based on their gene expression patterns across the entire model (Fig. 2A). The expression of all five transcripts, including RP5-875H18.9, RP11-1094H24.4, LINC01583, RP11-462L8.1, and LINC00973, was virtually undetectable in healthy iPSCs, healthy NPCs (20 and 40 days), and normal organoids. However, these markers were specifically and significantly induced in p53-KO NPCs (day 40), p53-KO organoids, and both U87 and U118 glioma cell lines. Among them, RP5-875H18.9 and RP11-1094H24.4 showed a restricted, tumor-specific pattern, with almost zero expression in all healthy samples. To our knowledge, this is the first report linking these two transcripts to glioma or glioblastoma. These five lncRNAs were selected as the top candidates of p53-deficient non-coding RNAs for further analysis.

**Fig. 2.**
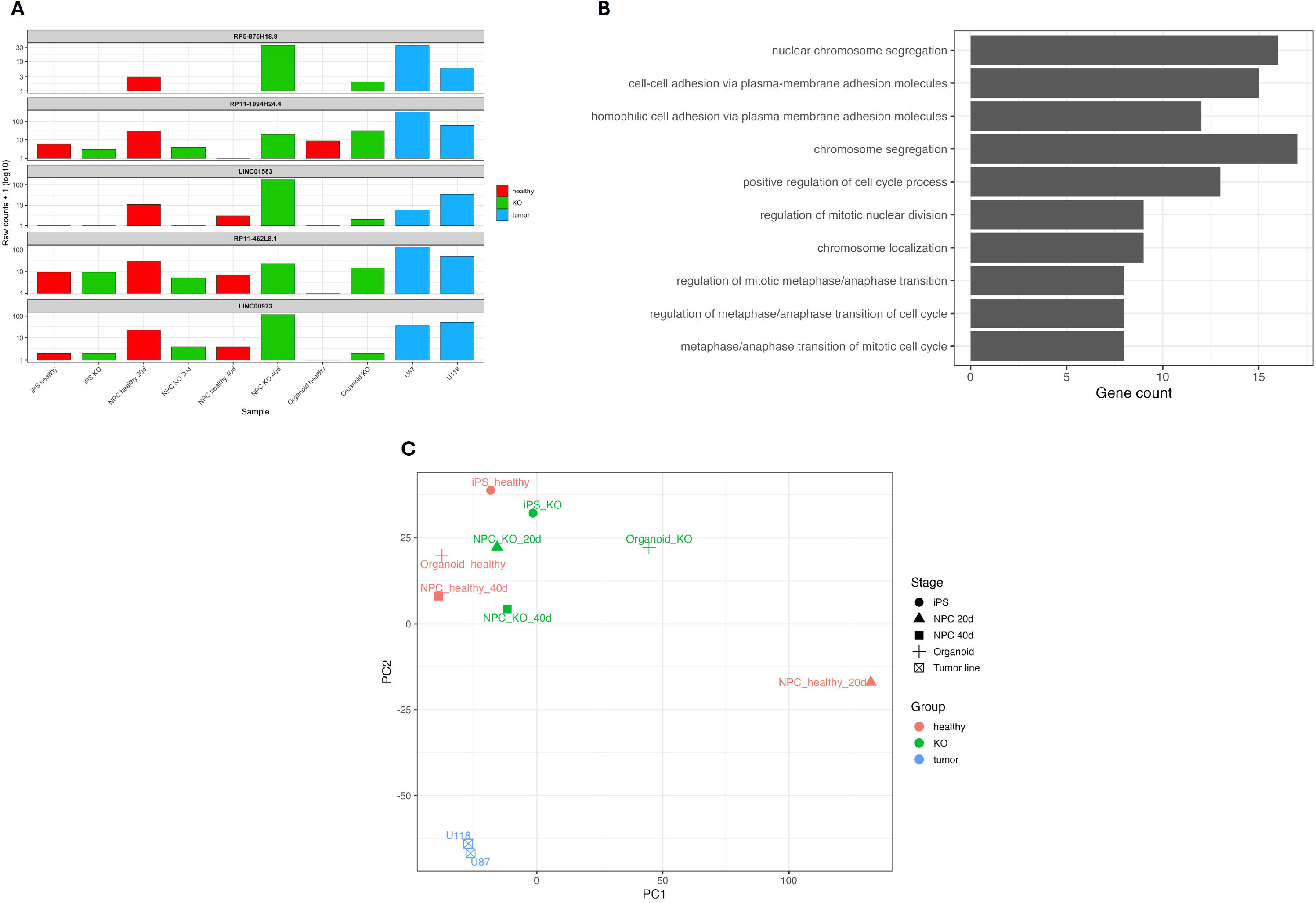
(A) Bar plots showing raw counts for the five key lncRNAs (RP5-875H18.9, RP11- 1094H24.4, LINC01583, RP11-462L8.1, and LINC00973) across all samples. All five transcripts are nearly undetectable in healthy iPSCs, healthy NPCs (20 d and 40 d), and healthy organoids, but are strongly expressed in late p53-KO stages (NPC 40 d and organoids) and in glioma cell lines U87 and U118. (B) Bar plot showing the top 10 Gene Ontology Biological Process terms enriched among genes upregulated in late p53-KO NPC (40 d), p53-KO organoids, and glioma lines. The dominant terms relate to nuclear chromosome segregation, positive regulation of the cell cycle process, and mitotic nuclear division. (C) Principal component analysis (PCA) was performed on the normalized expression levels of the five prioritized lncRNAs. Points are color-coded by group (healthy = red, p53-KO = green, tumor lines = blue) and shaped according to differentiation stage. Healthy iPSCs and early NPCs cluster on the left, while late p53-KO samples and glioma lines form a distinct tumor-like cluster on the right.

### 3.3. The tumor-like program is enriched for chromosome segregation and mitotic processes

Gene Ontology analysis of upregulated genes in late p53-KO and tumor-like samples showed significant enrichment in nuclear chromosome segregation, chromosome segregation, increased cell cycle activity, and mitotic nuclear division (Fig. 2B). This mitotic reprogramming module is typical of p53-deficient gliomagenesis and offers a biological basis for later heatmap analyses focusing on cell cycle genes.

### 3.4. p53 loss induces a progressive shift toward a tumor-like transcriptional state

To simulate early gliomagenesis, human iPSCs (control vs. p53 deletion) were differentiated into neural progenitor cells (NPCs) at 20 and 40 days, then further into tumor organoids. Additionally, the glioma cell lines U87 and U118 were used as reference samples showing tumor-like phenotypes. Principal component analysis (PCA) using the five prioritized lncRNAs clearly distinguishes the samples along a transition from healthy to malignant states (Fig. 2C). Late-stage (day 40) p53-KO NPCs, p53-KO organoids, and both glioma lines formed a distinct tumor-like cluster on the right side of the graph, while healthy iPSCs and early-stage NPCs clustered on the left. These results show that the absence of p53 alone causes a shift in the lncRNA profile that mimics the transcriptional state of established glioblastoma cells.

### 3.5. A late-stage “tumor-like program” links mitotic cell-cycle activation with specific lncRNAs

A heatmap of genes increasingly induced in late p53-KO and tumor-like phenotype samples (p53-KO 40d NPC, p53-KO organoids, U87, and U118) revealed a comprehensive “late KO/tumor-like molecular program” that included functional clusters related to immunity/inflammation, adhesion/membrane, secretion/signaling, stemness/transition, and associated lncRNAs (Fig. 3A). Most of these genes showed little to no expression in healthy iPSC, NPC (day 20 and 40), and organoids, but were consistently and strongly upregulated after p53 deletion.

**Fig. 3.**
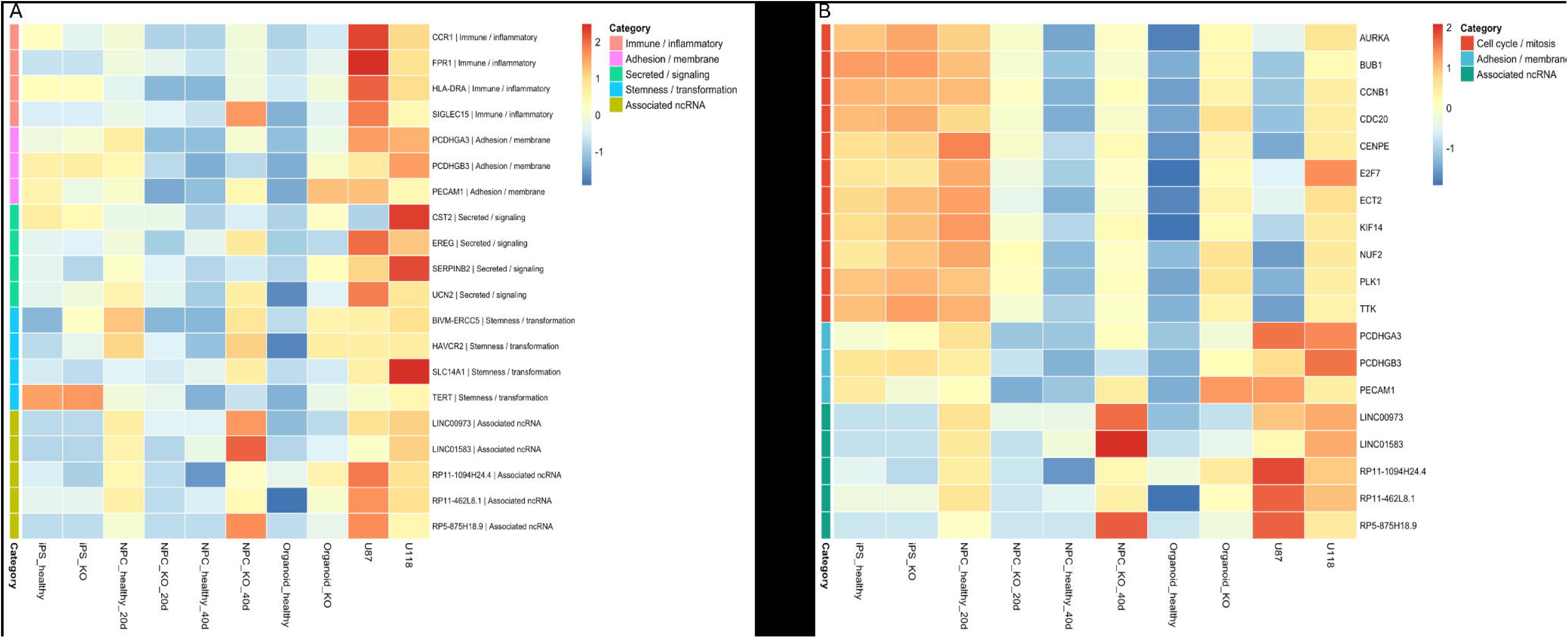
(A) Z-score normalized expression heatmap of genes strongly upregulated in late p53- KO stages and tumor-like samples (p53-KO 40 d NPC, p53-KO organoid, U87, and U118). Genes are categorized as immune/inflammatory (red), adhesion/membrane (pink), secreted/signaling (green), stemness/transformation (blue), or associated ncRNAs (yellow/green). Samples are ordered by differentiation stage, revealing coordinated activation specifically after p53 loss. (B) Focused heatmap highlighting the mitotic cell-cycle module (red) along with the five prioritized lncRNAs (green). Perfect co-regulation is observed specifically in late p53-KO stages and tumor lines, while healthy organoids remain at basal levels. The color scale indicates row-wise z-scores in both panels.

To explore the key drivers of this program, we next focused on the mitotic cell cycle module. A heatmap combining cell cycle/mitosis genes with five prioritized lncRNAs demonstrated complete co-regulation (Fig. 3B). Mitotic genes (e.g., AURKA, BUB1, CCNB1, CDC20, PLK1, TTK) and the five lncRNAs were not expressed in early stages of healthy samples but were significantly upregulated specifically in late p53-KO stage and tumor lines. Healthy organoids remained at basal levels, confirming that the absence of p53 is both necessary and sufficient to initiate a coordinated tumor-like mitotic signature.

### 3.6. Validation in primary patient tumors

To understand the clinical relevance of the findings, the TCGA glioma RNA-seq dataset was analyzed for the prioritized lncRNAs. LINC00973 and LINC01583 showed significant upregulation in glioblastoma (GBM) compared to low-grade gliomas (LGG) (Fig. 4A). Additionally, while LINC00973 showed significantly higher median expression in GBM, LINC01583 exhibited both high expression and high variability. Furthermore, survival analysis in the TCGA-GBM cohort showed that high LINC01583 expression was associated with significantly poorer overall survival (log-rank p = 0.0305), whereas LINC00973 showed a similar but non-significant trend (log-rank p = 0.283) (Fig. 4B). These results suggest that the tumor-specific lncRNA signature identified in the p53-KO organoid model is highly conserved in primary samples from patients with high-grade glioblastoma.

**Fig. 4.**
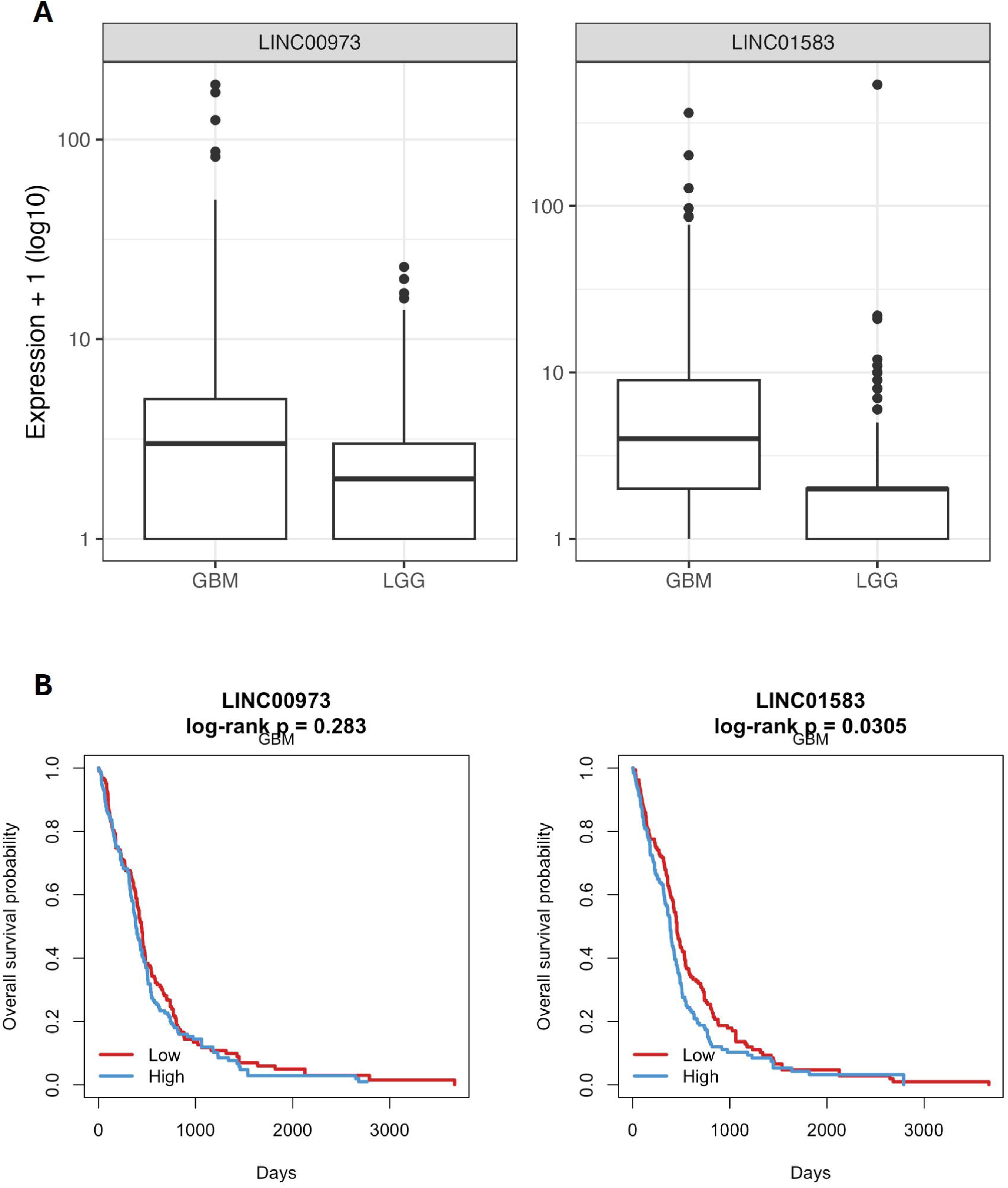
(A) Box plots of expression for LINC00973 and LINC01583 in TCGA glioblastoma (GBM) versus lower-grade glioma (LGG) cohorts. Both lncRNAs show significantly higher median expression and greater variability in GBM compared with LGG, confirming the clinical relevance of the tumor-specific signature identified in the p53-KO model. (B) Kaplan–Meier overall survival curves for GBM patients stratified by high vs. low expression of LINC00973 (left) and LINC01583 (right). High expression of LINC01583 was significantly associated with poorer overall survival (log-rank p = 0.0305), while LINC00973 showed a similar but non-significant trend (log-rank p = 0.283). Red = low expression, blue = high expression.

## 4. Discussion

This study, using an iPSC-p53KO organoid model, shows that p53 deficiency reorganizes lncRNAs into a late-emerging, tumor-like transcriptional network involved in mitotic chromosome instability. Our findings indicate that this molecular signature is conserved in primary glioblastoma patients, underscoring the significance of this pathway in disease development. Among the identified candidates, two lncRNAs, RP5-875H18.9 and RP11- 1094H24.4, show strong potential as early markers of gliomagenesis due to their highly specific expression in tumor tissue. These two transcripts have not been reported previously in glioma or glioblastoma, making them novel candidates for future research.

p53 is a gatekeeper in neural lineage differentiation, regulating self-renewal and proliferation, and promoting lineage commitment [9]. Loss of p53 shifts neural progenitor populations into a transient developmental suspension, which fails to exit the proliferative phase [10] and continues cell division [11]. This is particularly pronounced in the late 40-day NPC stage and in tumor organoids, indicating a delayed rather than an immediate consequence of p53 deficiency.

In this late tumor-like program, the five prioritized lncRNAs are integrated into cell cycle pathways and chromosomal segregation processes through co-regulation with mitotic regulators (AURKA, BUB1, CCNB1, and PLK1), suggesting they appear in the late stage to reinforce and stabilize the proliferative and progenitor-like state rather than initiating it. Additionally, RT-qPCR results supported this lineage-dependent reorganization. The p53-KO neural progenitors consistently maintained oncogenic and immature phenotypes (PAX6 and EGFR), while p53-KO organoids paradoxically retained neural differentiation markers (TUBB3 and MAP2). These findings show that p53 deficiency creates conditions for synergy between the prioritized lncRNAs and the mitotic machinery, leading to a “transcriptional lock” in cells in a glioblastoma-like transcriptional state.

Alongside the coding-gene program, we identified a distinct non-coding layer associated with the late p53-deficient state. The five prioritized lncRNAs (LINC00973, LINC01583, RP11- 1094H24.4, RP11-462L8.1, and RP5-875H18.9) were strongly enriched specifically in late p53-KO stages and tumor-like samples, indicating that lncRNA dysregulation accompanies rather than precedes the mitotic coding shift. Notably, RP5-875H18.9 and RP11-1094H24.4 have not previously been reported in glioma or glioblastoma, making this the first demonstration of their association with p53-loss-driven reprogramming. LINC00973 and LINC01583 were independently validated in TCGA glioma cohorts, showing significantly higher expression in primary GBM than in lower-grade glioma. Moreover, high LINC01583 expression was significantly associated with worse overall survival in TCGA-GBM patients (log-rank p = 0.0305), further strengthening its potential as a clinically relevant marker. LINC00973 showed a consistent but non-significant trend toward poorer survival. These findings strengthen the conclusion that at least part of the non-coding layer identified here reflects biologically relevant features detectable in glioblastoma.

Our research identifies a late p53-loss-associated transcriptional pathway during human brain development that includes both coding and non-coding transcripts. p53 deficiency causes a stage-specific, context-dependent lineage shift, with p53-KO organoids retaining more prominent neuronal differentiation markers. In contrast, late p53-KO NPCs show higher progenitor-biased and tumor-associated characteristics. The current analysis provides a solid, controlled platform for further mechanistic research, even though it does not establish functional causation for the prioritized lncRNAs. Specifically, the two transcripts (RP5- 875H18.9 and RP11-1094H24.4) that have not yet been identified in gliomas show promise as early biomarkers or as targets for RNA-based therapies.

To determine whether these lncRNAs actively promote or maintain the glioblastoma-like state, future studies using CRISPR interference, antisense oligonucleotides, and in vivo orthotopic models will be essential. This could lead to new reprogramming-focused strategies in neuro-oncology.

## Declarations

### Ethics statement

All experiments involving human iPSC-derived cells and organoids were conducted in accordance with institutional guidelines. Publicly available TCGA datasets were accessed under standard data-use agreements. No additional ethical approval was required for analysis of these publicly available datasets.

### Declaration of competing interest

The authors declare that they have no known competing financial interests or personal relationships that could have appeared to influence the work reported in this paper.

### Funding

*This work was supported by HUB Scienze della Vita della Regione Puglia - Codice Locale Progetto: T4-AN-01 (Cup: J73C22000470003)*.

### CRediT author contribution statement

- Fabio Stasi, Ph.D.: Conceptualization, Methodology, Investigation, Formal analysis, Data curation, Visualization, Writing – original draft, Writing – review & editing.
- Alireza Rajabzadeh Manzari, Ph.D.: Conceptualization, Methodology, Formal analysis, Supervision, Writing – review & editing.
- Velia Siciliano: Conceptualization, Supervision, Writing – review & editing.
- Elena Binda: Conceptualization, Supervision, Writing – review & editing.
- Valerio Pazienza: Conceptualization, Supervision, Writing – review & editing.
- Virgilio Brunetti: Conceptualization, Supervision, Writing – review & editing.
- Massimo De Vittorio: Funding acquisition, Supervision, Project administration, Writing – review & editing.

### Data availability

RNA-seq data generated in this study will be deposited in a public repository (e.g., GEO). The accession number will be provided upon acceptance. TCGA data are publicly available via the Genomic Data Commons portal (https://portal.gdc.cancer.gov).

## Acknowledgments

P.I. Elena Binda is supported by Italian Association for Cancer Research (AIRC) under IG 2024 - ID. 30722 project

## Abbreviations

GBM: glioblastoma multiforme
iPSC: induced pluripotent stem cell
lncRNA: long non-coding RNA
LGG: low-grade glioma
NPC: neural progenitor cell
PCA: principal component analysis
RNA-seq: RNA sequencing
TME: tumor microenvironment

## Notes

### Competing Interest Statement

The authors have declared no competing interest.

