## Supplementary material for "p53 knockout drives stage-specific long non-coding RNA reprogramming during human neural differentiation and glioma organoid tumorigenesis": Fig. S1

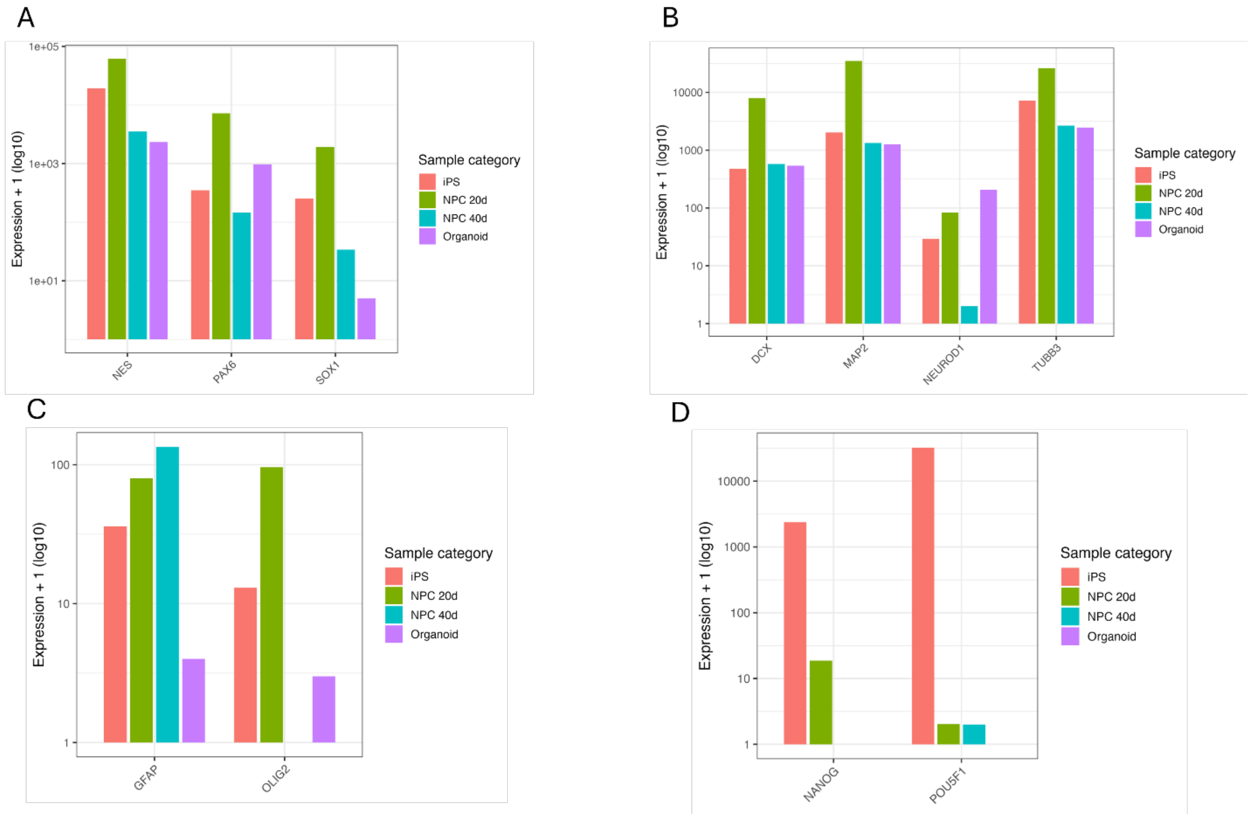


**Figure S1**. RNA-seq confirmation of successful neural differentiation in healthy samples. Bar plots of normalized expression ($\log_{10}$ scale + 1) for selected marker genes across healthy iPSC, NPC (20 d and 40 d), and cerebral organoid samples. (A) Neural progenitor markers (NES, PAX6, SOX1) peak during NPC stages. (B) Neuronal differentiation markers (DCX, MAP2, NEUROD1, TUBB3) are strongly upregulated in organoids. (C) Lineage markers (GFAP, OLIG2) appear in organoids, consistent with glial differentiation. (D) Pluripotency markers (NANOG, POU5F1) are high in iPSCs and rapidly downregulated upon differentiation. These patterns confirm that the healthy arm of the model successfully progresses through neural progenitor and organoid stages.
